# Draft genome sequence of the pulse crop blackgram [*Vigna mungo* (L.) Hepper] reveals potential R-genes

**DOI:** 10.1101/2020.06.21.163923

**Authors:** J Souframanien, Avi Raizada, Punniyamoorthy Dhanasekar, Penna Suprasanna

**Affiliations:** Nuclear Agriculture and Biotechnology Division, BARC, Trombay, Mumbai-400085, India; Homi Bhabha National Institute, Training School Complex, Anushaktinagar, Mumbai-400094, India

**Keywords:** Blackgram, *Vigna mungo*, Genome sequence, Transposons, SSRs, R-genes, next-generation sequencing

## Abstract

Blackgram [*Vigna mungo* (L.) Hepper] (2n = 2x = 22), an important Asiatic legume crop, is a major source of dietary protein for the predominantly vegetarian population. Here we construct a draft genome sequence of blackgram, for the first time, by employing hybrid genome assembly with Illumina reads and third generation Oxford Nanopore sequencing technology. The final *de novo* whole genome of blackgram is ~ 475 Mb (82 % of the genome) and has maximum scaffold length of 6.3 Mb with scaffold N50 of1.42 Mb. Genome analysis identified 18655 genes with mean coding sequence length of 970bp. Around 96.7 % of predicted genes were annotated. Nearly half of the assembled sequence is composed of repetitive elements with retrotransposons as major (47.3% of genome) transposable elements, whereas, DNA transposons made up only 2.29% of the genome. A total of 166014 SSRs, including 65180 compound SSRs, were identified and primer pairs for 34816 SSRs were designed. Out of the 18665 proteins, 678 proteins showed presence of R-gene related domains. KIN class was found in majority of the proteins (372) followed by RLK (79) and N (79). The genome sequence of blackgram will facilitate identification of agronomically important genes and accelerate the genetic improvement of blackgram.

## Introduction

Blackgram [*Vigna mungo* (L.) Hepper] is an annual leguminous crop belonging to family *Fabaceae* and sub-family *Papilionaceae*. This crop is a member of *Vigna* Savi (subgenus *Ceratotropis*), a genus belonging to the tribe phaseoleae, that includes other economically important grain legumes like cowpea (*Vigna unguiculata*(*L*.) Walp), mungbean (*Vigna radiata*(*L*.) R. Wilczek), common bean (*Phaseolus vulgaris* L.), pigeonpea (*Cajanus cajan* (L.) Millsp.) and adzuki bean (*Vigna angularis* (Willd.) Ohwi & Ohashi). Blackgram is a self pollinated diploid legume (2n =2x= 22) with genome size estimated to be 0.59 pg/1C (574 Mbp)^1^. It is popularly known as ‘urd bean’, ‘urd’ or ‘mash’ and is an excellent source of easily digestible good quality protein (25-26%), carbohydrates (60%), fat (1.5%), minerals, amino-acids and vitamins. In addition to being an important source of human food and animal feed, it also plays a significant role in sustaining soil fertility by improving soil physical properties and fixing atmospheric nitrogen. As a hardy legume tolerant to drought, blackgram is suitable for dry land farming and is predominantly grown as an intercrop or as a sole crop under residual moisture conditions post rice harvest. Blackgram originated in India and has been domesticated from its wild ancestral form *V. mungo* var. *silvestris*^2^. Extensively grown in south and south-east Asia from ancient times, it is one of the most highly prized pulses of India. India is the largest producer of blackgram, where about 5.0 million hectares are cultivated with an annual production of 3.8 million tonnes^3^.

In spite of its economic importance and surging demand for improved blackgram varieties, susceptibility to multiple pathogens, including mungbean yellow mosaic virus, powdery mildew, *Cercospora* leaf spot and leaf crinkle virus hinders cultivation and reduces produce yield and quality. In this regard, plant breeders and researchers are in the race for studying plant disease resistance mechanisms and identifying genes to develop varieties with durable resistance. Pyramiding of plant resistance genes in new cultivars is the most effective and environment friendly approach for plant disease control and reduction of yield losses. Plant disease resistance genes (R-genes) play a key role in recognizing proteins expressed by specific avirulence (Avr) genes of pathogens^4^. The proteins encoded by the resistance genes share common domains such as coiled-coil (CC), nucleotide binding region (NB), Toll-interleukin region (TIR), leucine rich region (LRR) and kinase domain (K). Hundreds of NBS-LRR, RLK and RLP genes have been reported in plants^5–8^, though such information is lacking in blackgram. This could be attributed to the lack of genomic resources coupled with limited understanding of the molecular basis of gene expression and phenotypic variation. Understanding the molecular basis of phenotypic variation and gene function is important for selective breeding traits such as increased yield, pest and disease resistance. Similarly, whole genome assemblies support GWAS studies to identify trait-specific loci and for genomic-based selective breeding^9^. To this end, whole-genome sequencing has been conducted on several commercial *Vigna* species such as mungbean, adzuki bean, cowpea, beach pea^5, 10–13^. Elucidation of the genome sequence of *V*. *mungo* var. *mungo* could reveal the general genome structure, repetitive sequence and R-gene composition of this legume species in comparison to closely related genomes and greatly assist comparative genomics with other well-studied legume genomes.

Next Generation Sequencing reads are too short to resolve abundant repeats in particular in plant genomes, leading to incomplete or ambiguous assemblies^14^. In the last few years, rapid innovations in sequencing technologies and improvement in assembly algorithms have enabled the creation of highly contiguous genomes. The development of third generation sequencing technologies that deliver long reads from single molecules and carry the necessary information to phase haplotypes over several kilobases have greatly improved the feasibility of *de novo* assemblies^15–17^. In view of the importance of the pulse crop in the Asiatic region and need for molecular detailing of trait based selection, we assembled a draft genome of *Vigna mungo* var. *mungo* using next generation platform Illumina paired end and mate pair reads combined with third generation Oxford Nanopore sequencing.

## Results

### Illumina and Nanopore sequencing of blackgram

We prepared three libraries for sequencing by Illumina HiSeqX Ten sequencer including 150 bp paired-end library and 5-7 kb and 7-10 kb mate-pair libraries. Whole genome sequencing using Illumina paired-end (PE) long insert generated 154,940,012 reads representing ~98x genome coverage. Sequencing of 2 mate-pairs of 5–7, and 7–10 kb yielded, 33,617,232 and 10,247,813 reads respectively, with an approximate coverage of 21.2xand 6.5xrespectively, and a grand total of 156 million mate-pair reads representing ~28xcoverage (Table S1). In addition, long read sequencing by Oxford Nanopore Sequencing Technology (ONT) was used to generate 1,633,898 long reads, having 10,425,220,236 bp and coverage of ~22x. A total of 11.5 Gb data was generated from whole genome library with an average read length of 6.4 kb and a maximum read length of 128.7 kb using Nanopore sequencer (Table S2). The complete genome was sequenced to a depth of ~148x, using both Illumina and ONT platforms.

### *De novo* assembly of blackgram genome and gene annotation

The raw reads generated from Illumina paired end, mate-pair and nanopore sequencing were processed and good quality reads were retained. Hybrid assembly was performed using Illumina and nanopore reads by MaSuRCA v3.3.4 hybrid Assembler. Scaffolds were further processed for super-scaffolds using PyScaf producing 1085 scaffolds with a N50 of 1.42 Mb (Table 1). Overall, the maximum scaffold assembled length was 6343.0 kb with median scaffold length of 67.9 kb. The total length of the produced scaffolds was 475 Mb (82 % of genome) for *Vigna mungo* cultivar Pant U-31.

**Table 1:** *De novo* assembly and annotation statistics of the blackgram genome.

|  |  |
| --- | --- |
| Scaffolds Generated : | 1085 |
| Maximum Scaffold Length (bp): | 63,43,804 |
| Minimum Scaffold Length (bp): | 510 |
| Average Scaffold Length (bp): | 438629 |
| Median Scaffold Length (bp): | 67909 |
| Total Scaffolds Length (bp): | 47,59,13,455 |
| Total Number of Non-ATGC Characters : | 3300 |
| Percentage of Non-ATGC Characters : | 0.001 |
| Scaffolds >= 100 bp : | 1085 |
| Scaffolds >= 200 bp : | 1085 |
| Scaffolds >= 500 bp : | 1085 |
| Scaffolds >= 1 Kbp : | 1048 |
| Scaffolds >= 10 Kbp : | 920 |
| Scaffolds >= 1 Mbp : | 168 |
| N50 value : | 14,26,686 |
| Number of genes | 18655 |
| Average gene length | 971 bp |
| Maximum gene length | 18.07 kb |
| Minimum gene length | 201 bp |
| Number of genes | 18655 |

The gene prediction and annotation of the assembled genome was carried out using AUGUSTUS software. In total 18655 genes were identified with average coding sequence length of 971 bp. The maximum and minimum sequence lengths were 18.07 kb and 201 bp, respectively (Table 1). A total of 18049 genes (96.75%) of predicted genes could be functionally annotated with gene ontology and pathway information (Table S3). Gene ontology provides a system to categorize description of gene products according to three ontologies: molecular function, biological process and cellular component. Of the 18049 annotated genes, majority (53.0 %)were assigned with molecular function, followed by cellular components (30.7 %) and biological functions (13.8 %). Among the assignment made to the molecular function, a large proportion of the sequences represented nucleic acid binding (13.7%) followed by ATP binding (10.8%) and zinc ion binding (8.4%). Among those with cellular component function, the majority represented integral component of membranes (18.2%) followed by nucleus (6.7%) and cytoplasm (1.6%). Under the biological process category, more sequences were assigned to DNA integration (6.9%) followed by regulation of transcription (1.9%) and carbohydrate metabolic process (1.4%) (Fig. 1). Pathway assignments were carried out according to the Kyoto Encyclopedia of Genes and Genomes (KEGG) pathway database. A total number of 4322 unique KEGG pathways were identified (Table S4), of which the majority of sequences were grouped into protein families metabolism (737) followed by carbohydrate metabolism (600). Orthologous gene comparison studies using genes from *Vigna mungo* (PantU-31), *Vigna radiata* and *Vigna angularis* were carried out using Ortho Venn13. A total of 7534 gene clusters were shared by all three species, while 446 gene clusters were specific to *Vigna mungo* (Fig. 2).

**Fig. 1.**
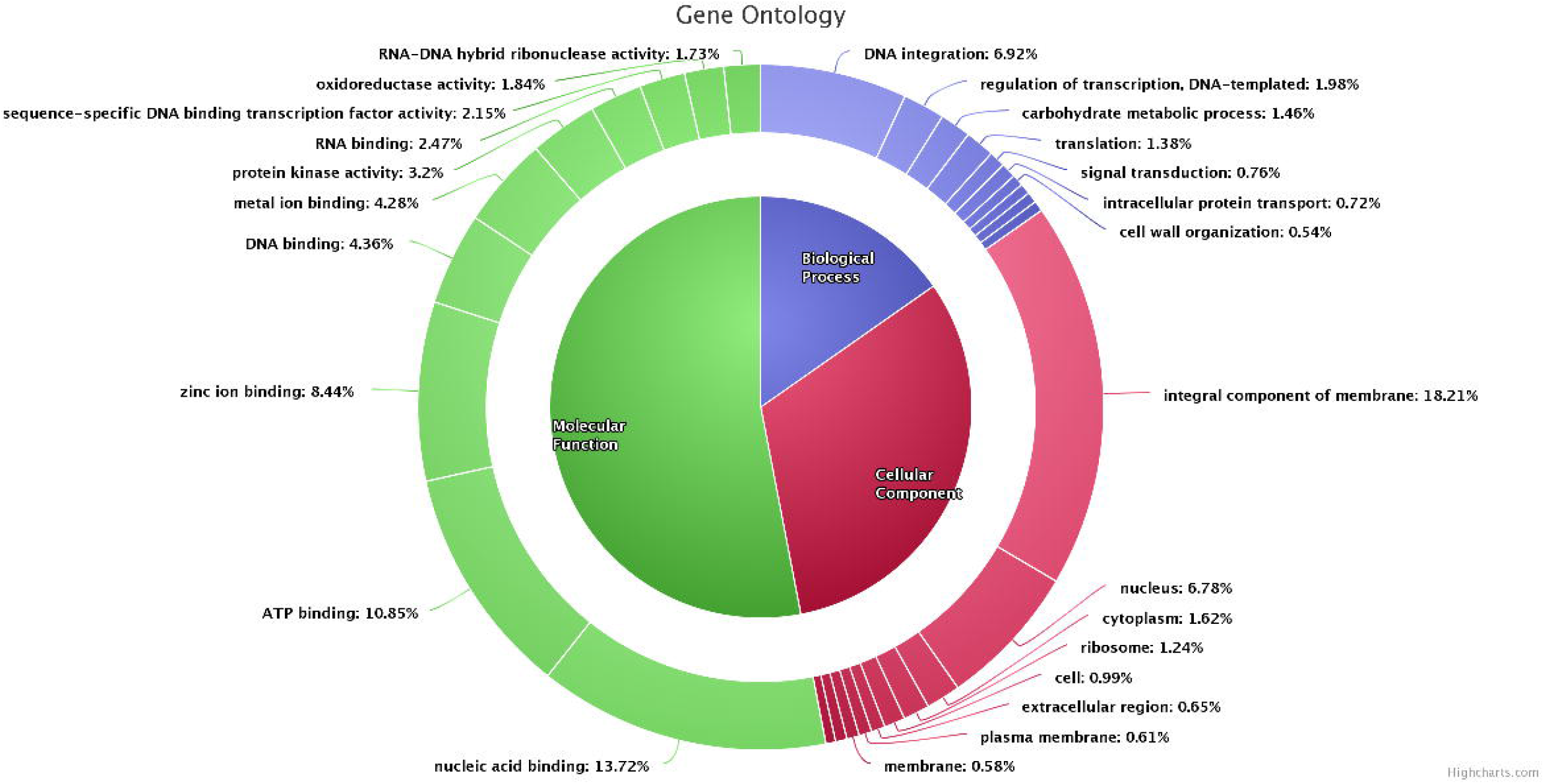
Gene Ontology chart of *Vigna mungo*.

**Fig. 2.**
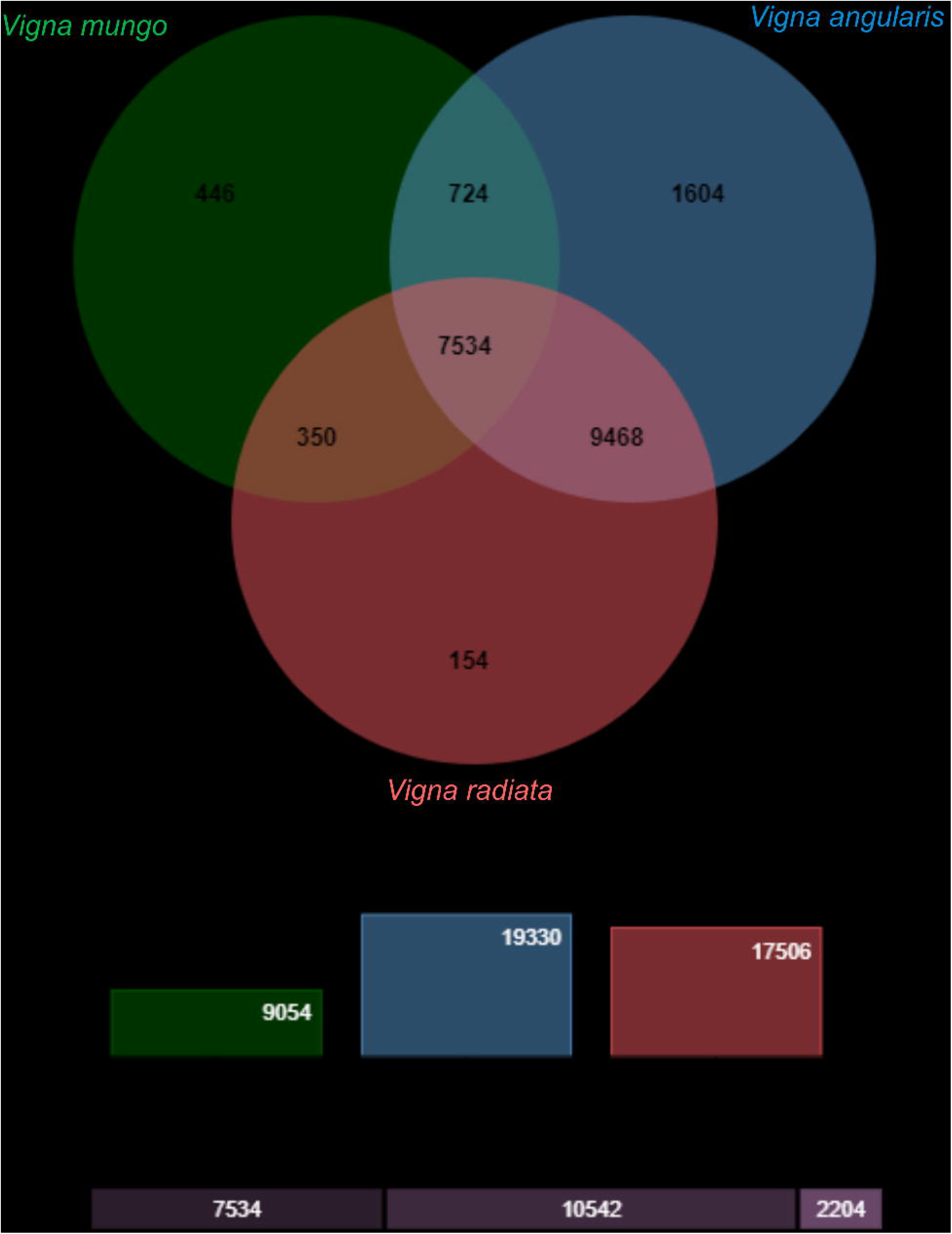
Venn diagram showing shared orthologous gene clusters among *V. mungo, V. radiataand V. angularis*.

### Prediction of transposons

The presence of transposons in the assembled genome was predicted using TREP (TRansposable Elements Platform). Repetitive sequences occupy 49.6% of the *V. mungo* genome as revealed by homology- and structure-based surveys. Majority of the transposable elements were retrotransposons (47.3% of genome), whereas DNA transposons made up only 2.29% of the genome (Table 2). Long terminal repeat (LTR) retrotransposons forming the predominant class of transposable elements in *V*. *mungo* genome showed homologies with that of *Metrosideros polymorpha, Blumeria graminis_tritici, Sorghum bicolor, Triticum aestivum, Hordeum vulgare, Brachypodium distachyon, Arabidopsis thaliana* and *Oryza sativa* genomes. Overall, 47.3% of the repetitive DNA was long terminal repeat retrotransposons of which 13.4 % were Gypsy type and 34.5% were Copia type elements. In contrast, class II DNA transposons, including Mutator, PIF-Harbinger, hAT, Helitron, and Tc1-Mariner, accounted for 2.3% of the blackgram genome. The rolling-circle Helitron (DHH) superfamily is relatively abundant at 1.3% of the genome (Table S5). Only 3.1% of the TE sequences were unclassified.

**Table 2.**
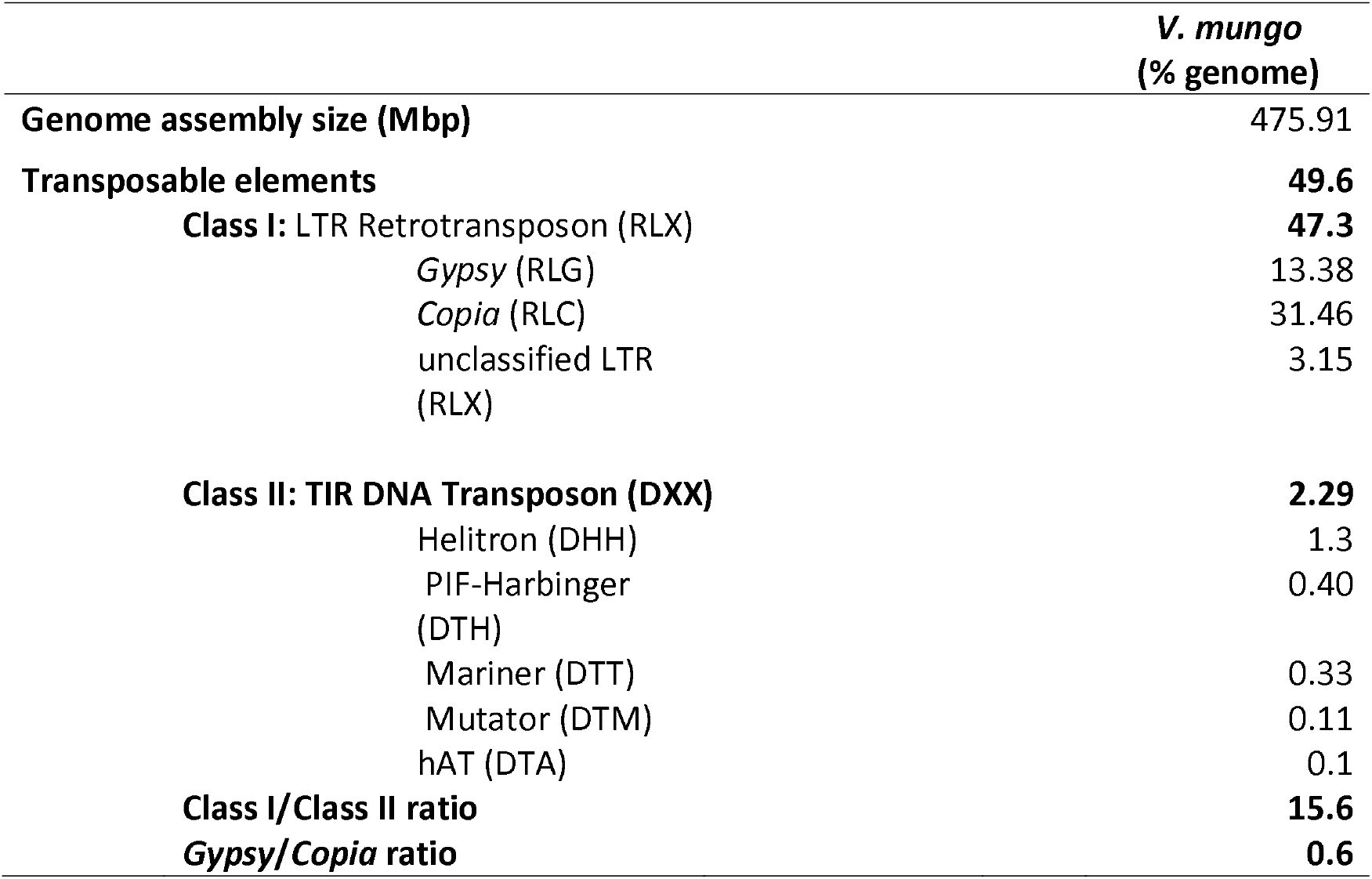
Annotated repeat abundances in blackgram. The major represented classes, super-families, and subgroups of transposable elements as determined by automated annotation and classified according to the scheme of Wicker et al. (2007), as well as other major repeat types are presented.

|  | ***V. mungo***<br>**(% genome)** |
| --- | --- |
| **Genome assembly size (Mbp)** | 475.91 |
| **Transposable elements** | **49.6** |
| **Class I: LTR Retrotransposon (RLX)** | **47.3** |
| *Gypsy* (RLG) | 13.38 |
| *Copia* (RLC) | 31.46 |
| unclassified LTR (RLX) | 3.15 |
| **Class II: TIR DNA Transposon (DXX)** | **2.29** |
| Helitron (DHH) | 1.3 |
| PIF-Harbinger (DTH) | 0.40 |
| Mariner (DTT) | 0.33 |
| Mutator (DTM) | 0.11 |
| hAT (DTA) | 0.1 |
| **Class I/Class II ratio** | **15.6** |
| ***Gypsy/Copia* ratio** | **0.6** |

### Simple Sequence Repeats (SSR) prediction

SSRs were detected using Microsatellite Identification Tool (MISA v1.0). A total of 166014 SSRs were identified from 989 scaffolds (Table 3). More than one SSR were present in 953 scaffolds and 65180 SSRs were of the compound type. SSR loci with di- and tri-nucleotides constituted 103955 (62.6%) of the identified loci. The proportions of di-, tri-, tetra-, penta-, and hexa-nucleotide repeats were 38.1%, 24.5%, 36.4%, 0.69%, and 0.24%, respectively (Table S6). The number of repeats varied from 6-61 for di-nucleotides, 5-361 for trinucleotides, 3-7 for tetra-nucleotides, 5-19 for penta-nucleotides and 5-14 for hexanucleotides. The most prevalent di-, tri-, tetra-, penta-, and hexa-nucleotide repeats were AT (22.6%), AAT (3.9%), TTTA (5.1%), AAAAT (4.6%) and ATGTTG (1.9%), respectively (TableS7). Of the 166014 SSR motifs identified, PCR primer pairs were successfully designed for 34816 SSR loci. Details about primer sequences and expected product sizes for 34816 SSR loci are provided in supplementary table (Table S7).

**Table 3.** Number and distribution of SSRs identified in the blackgram (*Vigna mungo*) cv. Pant U-31 genome.

| **Description** | ***V. mungo* genome** |
| --- | --- |
| Total number of sequences examined | 1085 |
| Total size of examined sequences (bp) | 475913455 |
| Total number of identified SSRs | 166014 |
| Number of SSR containing sequences | 989 |
| Number of sequences containing more than 1 SSR | 953 |
| Number of compound SSRs (i.e c) | 65180 |
| p2 | 63220 |
| p3 | 40735 |
| p4 | 60512 |
| p5 | 1146 |
| p6 | 402 |

### Identification of disease resistance genes

A total of 18665 protein sequences were analysed for resistance (R) genes related domains and motifs with the help of DRAGO 2 (Disease Resistance Analysis and Gene Orthology) pipeline of Plant resistance gene database (PRGDB). Out of 18665 proteins, 678 proteins showed presence of R-gene related domains. Majority of the proteins (269 proteins) contained TM-kinase domains and kinase formed the major class (372 proteins) (Table S8). One hundred and six proteins (15.6 %) were found to have Nucleotide Binding Sites (NBS) (Table 4). A total of 156 proteins showed a single type domain (112 kinase, 19 LRR, 15 NBS, 6 TIR, 3 TM, and 1 CC) (Table 4). While remaining proteins harboured more than one domain types such as NBS-TM, TIR-NBS etc. The LRR-TM-Kinase-CC, NBS-LRR-TM, NBS-CC-TM-TIR and NBS-TM-TIR domain combinations were found in 2, 2, 1 and 1 proteins, respectively. Among the different classic R-gene classes majority were found to be of kinases (KIN)(55.0%) followed by transmembrane receptors (RLP or RLK)(15.5%) and no proteins were found to represent the class of cytoplasmic proteins (CNL and TNL). The classic R-gene classes RLP (Ser/Thr-LRR) and RLK (Kin-LLR) were found in 26 and 79 proteins, respectively. R-domain occurrence in the full dataset showed that the NBS and LRR domains were found in 6 classes each, followed by the KIN domain in 5, and TIR domains in 3 classes. Likewise, proteins showing other classes such as TN, TRAN, NL, CNK, C, CNT and CLK were found in 3, 2, 2, 1, 1, 1 and 1 proteins, respectively (Table 4). Seventy-one R-genes were identified based on their homologies with mungbean, cowpea and adzuki bean sequences (Table S9).

**Table 4.** Prediction of Resistance genes domains/motifs present in proteins identified from whole genome sequencing of blackgram cultivar PantU-31 with the help of DRAGO pipeline of Plant resistance gene database

| **Domain/motif Types** | **Number of Proteins** | **Class** | **Number of Proteins** |
| --- | --- | --- | --- |
| Kinase | 112 | KIN | 373 |
| TM-Kinase-LRR | 75 | RLK | 79 |
| TM-Kinase | 269 | CK | 44 |
| CC-TM-Kinase | 33 | L | 19 |
| LRR | 19 | N | 79 |
| NBS-TM | 61 | TRAN | 2 |
| TM | 3 | CNK | 1 |
| CC-Kinase | 11 | CN | 24 |
| NBS-CC | 7 | TN | 3 |
| TIR-NBS | 2 | RLP | 26 |
| LRR-Kinase | 4 | T | 10 |
| LRR-TM | 26 | NK | 6 |
| TIR | 6 | CL | 7 |
| NBS-CC-TM | 17 | C | 1 |
| NBS | 15 | NL | 2 |
| CC-LRR | 6 | CNT | 1 |
| LRR-TM-Kinase-CC | 2 | CLK | 1 |
| CC | 1 | Total no: of proteins | 678 |
| TM-TIR | 4 |  |  |
| CC-LRR-TM | 1 |  |  |
| NBS-LRR-TM | 2 |  |  |
| NBS-CC-TM-TIR | 1 |  |  |
| NBS-TM-TIR | 1 |  |  |
| Total no: of proteins | 678 |  |  |

## Discussion

A better understanding of blackgram genetics is crucial for more efficient breeding in light of an anticipated increase in biotic and abiotic stresses that may accompany climate change. Whole-genome sequences are an important resource for evolutionary geneticists studying plant domestication, as well as breeders aiming to improve crop varieties. We sequenced *V*. *mungo* using Illumina PE and Nanopore with a coverage of 148x. In the present study, we have developed an integrated approach, including next generation and Nanopore sequencing and assembled genome using MaSuRCA hybrid assembler. The final assembly comprised of 1085 scaffolds (N50=1.43 Mb). Hybrid assembly through combinational sequencing is a useful approach in obtaining accurate sequence data. Moreover, the production of long-reads while using third generation sequencing (Nanopore) overcomes the weakness of assembling short-reads by minimizing the generation of gaps or covering the repetitive sequences that appear in the plant genomes. In addition, while only considering the accuracy, short-reads can be used for error-correction by aligning them to long-reads, which enable the increased accuracy of the genome assembly^18^. We constructed 475 Mb (82%) of the total estimated *V. mungo* var. *mungo* genome and identified 18,655 protein-coding genes and 446 *Vigna* mungo gene clusters. This is the first draft genome sequence of *Vigna mungo*. The assembly generated will also advance comparative genomics in *Vigna* species, as whole genome sequences of prominent *Vigna* species including mung bean, adzuki bean and cowpea are already available^5,11,12^. Of the 18,655 predicted genes, 18049 could be functionally annotated. In *V*. *radiata* genome, 22,427 genes were annotated with high confidence^5^. Most of the gene annotations were comparable to the annotation of immature seed transcriptome sequence of blackgram^19^. Orthologous gene comparison studies using genes from *Vigna mungo* (PantU-31), *Vigna radiata* and *Vigna angularis* was carried out using Ortho Venn13. There was a total of 7534 gene clusters shared by all three species, and 446 clusters were composed of only *Vigna mungo* specific proteins. High degree conservation and collinearity between blackgram and adzuki bean was revealed through comparative mapping^20^. Gene order conservation between closely related legume species (*V*. *angularis* var. *angularis*, *V*. *radiata* var. *radiata*, and *P*. *vulgaris*) has been exploited in synteny based scaffolding approach in genome assembly^11^. Similarly, Cowpea chromosomes Vu02, Vu03 and Vu08 also have one-to one relationship with the other two *Vigna* species (mungbean and common bean) suggesting that these chromosome rearrangements are characteristic of the divergence of *Vigna* from *Phaseolus*^12^.

### Transposable elements (TEs)

In plants, transposable elements are a major driver of genome expansion. Retrotransposons are the predominant TEs in large plant genomes and are further divided into class I, those flanked by long terminal repeats (LTRs) and those devoid of them. The class II elements, on the other hand, transpose via DNA intermediate and possess terminal inverted repeats (TIRs), which serves as sites of excision and re-integration by element-encoded transposase^21^. Homology and structure based analysis revealed that LTRs are the predominant class of transposable elements in the *Vigna mungo* genome, consistent with other legume species^5,22–26^. Of the long terminal repeat (LTR) retrotransposons, elements of the *Copia* superfamily^27^ (code RLC) are 0.6 times more abundant than *Gypsy* (code RLC) elements in blackgram. However, *Gypsy* element was found to be more abundant in the related *Vigna* species such as mungbean, adzuki bean and cowpea^5,11,12^. The DNA, or class II, transposons comprise2.3% of the genome, with Mutator, PIF-Harbinger, hAT, Helitron, and Tc1-Marinerbeing the major groups of classical ‘cut-and-paste’ transposons in blackgram. The rolling-circle Helitron (DHH) superfamily relatively abundant in blackram is in consistent with cowpea^12^.

TEs are potential reservoirs of phenotypic variation and phenotypic plasticity^28^. Moreover, TEs can directly assist the crop improvement programs through molecular marker approach. The presence of TEs, often close to or within the stress responsive quantitative trait loci (QTLs), especially plant defense genes, along with the traditional attributes of a molecular marker, make them the markers of choice for diversity studies and trait mapping^29,30^. While more studies would be necessary to understand the functional effects of these insertions, long-read sequences have greatly improved the assembly and identification of repeat types.

### Simple Sequence Repeats

The development of genomic resources is critical for crop improvement programmes. NGS has allowed the discovery of a large number of DNA polymorphisms, such as SNP and InDels markers, in a relatively short time at low cost^31^. Among 166014 SSRs (excluding mono nucleotide repeats) identified, the proportions of dinucleotide repeats were higher (38.1%) compared to other repeats in *V*. *mungo*. Similarly, dinucleotide repeats were found to be higher (71.3%) compared to other repeats in *V*. *radiata*^5^. Proportion of tri-, tetra-, penta-, and hexa-nucleotide SSRs were more or less same in comparison to *V*. *radiata* (24.6%, 2.5%, 1.2%, 0.2%) and lower than *V*. *marina* (49%, 3%, 7%, 5%) except for tetra-nucleotide repeats. Tetra-nucleotide repeats in *V*. *mungo* were found to be higher (36.4%) in comparison to *V. radiata* (2.5%) and *V. marina* (3.0%). However, the number of compound SSRs was higher (39.2%) than that in *V. radiata* (35.9%) and *V. marina* (10.08%)^5,13^. These findings indicate that the genome of *V. mungo* is more complex than that of *V*. *radiata*. To date, few efforts have been made to develop sufficient genomic resources in *Vigna*. This first genome sequencing effort in *V*. *mungo* has generated SSRs and functional annotations for a huge set of genes. This information holds great promise for use in trait mapping, genomic selections and diversity assessment.

### Disease resistance genes

Whole genome sequencing has enabled genome-level investigation of the R-gene family in crop plants such as mungbean, chickpea, rice, tomato5-8. In blackgram 3.6 % of the total genes were found to contain R-genes which is higher (1.2%) than that reported for *Medicago*^32^ and lower (5.27%) than that reported for *Arabidopsis^33^*. Plants possess a sophisticated immune system based on their ability to recognize phytopathogens. The activation of this system is based on the presence of specific receptors encoded by R-genes. Resistance genes are grouped as either nucleotide binding site leucine rich repeat (NBS-LRR) or transmembrane leucine rich repeat (TM-LRR)^34^. NBS-LRR proteins encoded by resistance (R) genes play an important role in pathogen recognition process and the activation of signal transduction in the response to pathogen attack. NBS-LRR can be further classified as toll/interleukin receptor (TIR)-NBS-LRR (TNL) or non-TNL/coiled coil-NBS-LRR (CNL)^34^. Both TNL and CNL specifically target pathogenic effector proteins inside the host cell, termed effector triggered immunity (ETI) response^35^. In *Vigna mungo* 15.6 % of total identified R-gene related sequence showed NBS domain. In *Vigna mungo* transmembrane leucine rich repeat (TM-LRR) class such as receptor like kinase (RLK) and receptor like protein (RLP) accounted for 15.5 % of the R-genes identified. RLPs and RLKs are pattern recognition receptors (PRRs) that mediate pathogen/microbe associated molecular pattern (PAMP/MAMP) triggered immunity (PTI/MTI) to allow recognition of a broad range of pathogens^35^. Development of diagnostic molecular markers associated with key disease resistance gene would aid in molecular resistance breeding.

In this study, the black gram genome was assembled using hybrid approach with the size of 475Mb. A total of 18655 genes were predicted from the assembled genome. Further, the predicted genes were annotated with gene ontology and pathway information. The presence of transposons and SSRs in the assembled genome was also predicted. Blackgram is grown mostly in developing countries and lack of genome sequence has delayed the implementation of molecular breeding in this *Vigna* species. The whole-genome sequence and SSR discovery will thus boost genomics-assisted selection for blackgram genetic improvement.

## Methods

### DNA Extraction

Blackgram (*V. mungo* var. *mungo*) cultivar PantU-31 was used for whole genome sequencing. DNA was extracted from 50-100 mg young leaves using Qiagen DNA easy Plant Mini kit following manufacturer’s instructions. Extracted genomic DNA was quantified and assessed for quality using Nanodrop2000 (Thermo Scientific, USA), Qubit (Thermo Scientific, USA) and agarose gel electrophoresis.

### Illumina library preparation and sequencing

Whole genome sequencing (WGS) libraries were prepared using Illumina-compatible NEXTFlex Rapid DNA sequencing Bundle (BIOO Scientific, Inc. U.S.A.) at Genotypic Technology Pvt. Ltd., Bangalore, India. Briefly, 300 ng of Qubit quantified DNA was sheared using Covaris S220 sonicator (Covaris, Inc. USA) to generate specific fragments in the size range of 300-400 bp. The fragment size distribution was verified on Agilent 2200TapeStation and subsequently purified using High prep magnetic beads (Magbio Genomics). Purified fragments were end-repaired, adenylated and ligated to Illumina multiplex barcode adaptors as per NEXTflex Rapid DNA sequencing bundle kit protocol.

### Matepair Illuminalibrary preparation

Mate pair sequencing library was prepared using Illumina-compatible Nextera Mate Pair Sample Preparation Kit (Illumina Inc., Austin, TX, U.S.A.). About 4 μg of genomic DNA was simultaneously fragmented and tagged with mate pair adapters in a transposon based tagmentation step. Tagmented DNA was then purified using AMPure XP magnetic beads (Beckman Coulter, U.S.A.) followed by strand displacement to fill gaps in the tagmented DNA. Strand displaced DNA was further purified with AMPure XP beads before size-selecting the fragments on low melting agarose gel. Size selected fragments were circularized in an overnight blunt-end intra-molecular ligation step that resulted in circular DNA with the insert flanked mate pair adapter junction. Circularized DNA was sheared using Covaris S220 sonicator (Covaris, Woburn, Massachusetts, USA) to generate fragment size distribution from 300 bp to 1000 bp. Sheared DNA was purified to collect the Mate pair junction positive fragments using Dynabeads M-280 Streptavidin magnetic beads (Thermo Fisher Scientific, Waltham, MA, U.S.A.). Purified fragments were end-repaired, adenylated and ligated to Illumina multiplex barcode adaptors as per Nextera Mate Pair Sample Preparation Kit protocol. Sequencing library, thus constructed, was quantified using Qubit fluorometer (Thermo Fisher Scientific, MA, USA) and its fragment size distribution was analyzed on Agilent 2200 TapeStation. The libraries were sequenced on Illumina HiSeq X Ten sequencer (Illumina, San Diego, USA) using 150 bp paired-end chemistry following manufacturer’s instructions.

### Nanopore library preparation and sequencing

A total of 1.5 μg of gDNA was end-repaired (NEBnext ultra II end repair kit, New England Biolabs, MA, USA) and purified using 1x AmPure beads (Beckmann Coulter, USA). Adapter ligation (AMX) was performed at RT (20 °C) for 20 minutes using NEB Quick T4 DNA Ligase (New England Biolabs, MA, USA). The reaction mixture was purified using 0.6X AmPure beads (Beckmann Coulter, USA) and sequencing library was eluted in 15 μl of elution buffer provided in the ligation sequencing kit (SQK-LSK109) from Oxford Nanopore Technology (ONT). Sequencing was performed on GridION X5 (Oxford Nanopore Technologies, Oxford, UK) using SpotON flow cell R9.4 (FLO-MIN106) in 48 hrs sequencing protocol on MinKNOW 2.1 v18.08.3with Albacore (v1.1.2)^36^ live base calling enabled with default parameters.

### Primary data analysis

The data obtained from the Illumina sequencing run was demultiplexed using Bcl2fastq softwarev2.20(https://sapac.support.illumina.com/sequencing/sequencing_software/bcl2fastqconverson-software.html)and FastQ files were generated based on the unique dual barcode sequences. The sequencing quality was assessed using FastQC v0.11.8 software^37^. The adapter sequences were trimmed using Trimgalore v0.4.0^38^ and bases above Q30 were considered and low quality bases were filtered off during read pre-processing and used for downstream analysis. Similarly, the Nanopore reads were processed with default settings using Porechop tool (https://github.com/rrwick/Porechop). The pre-processing of Nanopore data retained 99.9 % of data.

### *De novo* Genome assembly and gene annotation

Hybrid assembly was performed using Illumina and nanopore processed reads by MaSuRCA v3.3.4 hybrid Assembler^39^ with standard parameters. The assembled contigs were utilized to generate larger scaffolds using pyScaf(v1) software (https://github.com/lpryszcz/pyScaf). The generated assembled genome of ~ 475MB size was used for further analysis. The gene prediction and annotation of the assembled genome was carried out using AUGUSTUS tool^40^. It helped in the identification of protein-coding genes and their exonic-intronic structure in the genome in order to improve the accuracy and completeness of the annotation. AUGUSTUS predicted proteins were checked for similarity against Uniprot Phaseoleae database^41^ using DIAMOND blastp^42^ program with an e-value of 1e-5 for gene ontology and annotation. Pathway analysis was performed using KAAS server^43^. KAAS (KEGG Automatic Annotation Server) provides functional annotations of genes in a genome by amino acid sequence comparisons against a manually curated set of ortholog groups in KEGG genes. Comparative analysis of the organization of orthologous gene clusters were carried out using genes of *Vigna mungo, Vigna radiata* and *Vigna angularis* by OrthoVenn^44^ with E-value of 0.01and inflation value of 1.5.

### Identification of Transposable elements and Simple Sequence Repeats (SSR’s)

Transposon elements analysis was performed against TREP (TRansposable Elements Platform)^45^ which is a curated database of transposable elements (TEs)(http://botserv2.uzh.ch/kelldata/trep-db/index.html). Each consensus representing a structural variant of a TE family was classified according to its structural and functional features. TEs classifications were based on its ability to replicate in a host genome using various transposition mechanisms and are divided into two classes based on their replication mechanism. Retrotransposons (class I) use an RNA intermediate for transposition while DNA transposons (class II) use a DNA intermediate for transposition^27^. The genome sequence was checked for homology with TREP database using BLASTn^46^ and the genomic positions having homology with known TEs were identified.

SSRs were identified from the genome sequence using Micro SAtellite identification tool (MISA) [http://pgrc.ipk-gatersleben.de/misa/]. This predicted polymorphic loci of 1-6bp length in nucleotide sequences. Repeats were identified in each scaffold sequences using MISA Perl script. In this study, the SSRs were considered to contain motifs with two to six nucleotides in size and a minimum of 6, 6, 3, 5, 5 contiguous repeat units for di-, tri-, tetra-, penta- and hexa-nucleotides, respectively. Mononucleotide repeats were not included in the SSR search criteria. Based on MISA results, primers were designed to SSR motifs using either with WebSat (http://purl.oclc.org/NET/websat/) online software^47^ or batch primer3 ver1.0^48^. For designing PCR primers, parameter for optimum primer length was 22 mer (range: 18–27 mer), optimum annealing temperature was 60 °C (range: 57–68 °C), GC content was 40–80%, and other parameter values as default.

### Identification of disease resistance genes

Disease Resistance Analysis and Gene Orthology (DRAGO v.2) pipeline was used to predict and annotate the disease resistance genes from the Plant Resistance Genes database (PRGdb 3.0; http://prgdb.org) with curated reference R-genes^49,50^. DRAGO was executed with peptide sequence file from *V*. *mungo* var. *mungo* as an input to define the normalization value and the minimum score thresholds. Specifically, the previously created 60 HMM (hidden Markov model) modules were used by DRAGO 2 to detect LRR, Kinase, NBS and TIR domains and compute the alignment score of the different hits based on a BLOSUM62 matrix. The normalization value was the absolute smallest similarity score found among the input sequences considering all domains. The minimum score thresholds were calculated from the smallest similarity score reported in a specific domain among the input sequences. DRAGO 2 generated files with numeric matrix that represented the similarity score of every single protein input to each HMM profile, the domain name, start position, end position, resistance class and identification for every putative plant resistance protein.

## DATA AVAILABILITY

The de novo genome assembly has been deposited at GenBank under submission ID, Bioproject PRJNA631562 and biosamplesSUB7425717.

## ACKNOWLEDGEMENT

Authors thank Dr. P. Venugopalan, Associate Director, Biosciences Group, Bhabha Atomic Research centre, Trombay, Mumbai, for his kind support and encouragement for execution of the project.

## AUTHOR CONTRIBUTIONS

JS conceived the idea, coordinated the sequencing and wrote the manuscript. AR contributed to R-gene analysis. PD contributed SSR analysis and primer designing. PS supervised the study. All authors have read and approved the manuscript.

