## Supplementary Table 1 for "Draft genome sequence of the pulse crop blackgram [*Vigna mungo* (L.) Hepper] reveals potential R-genes": Table S1.docx

**Table S1:** Raw data statistics of blackgram genome reads generated by Illumina HiSeq and ONT.

| **Sample** | **Platform** | **Library and chemistry** | **No. of raw reads** | **No. of processed reads** | **Coverage** |
| --- | --- | --- | --- | --- | --- |
| SO_8668_PE | HiSeq | PE (150 x 2) | 154940012 | 140116780 | 98x |
| SO_8668_MP_5-7 kb | HiSeq | MP (150 x 2) | 33617232 | 26742270 | 21x |
| SO_8668_MP_7-10 kb | HiSeq | MP (150 x 2) | 10247813 | 7586306 | 6.5x |
| SO_8668_NP | ONT | Long read | 1633898 | 1633786 | 22x |

Abbreviations: kb, kilobases; PE, paired-end; MP, mate-pair; ONT, Oxford Nanopore Technology.
