## Supplementary Table 2 for "Draft genome sequence of the pulse crop blackgram [*Vigna mungo* (L.) Hepper] reveals potential R-genes": Table S2.docx

**Table S2:** Raw data statistics of blackgram genome reads generated by ONT.

| **Data type** | **Raw data** | **Processed** |
| --- | --- | --- |
| Reads Generated | 1633898 | 1633786 |
| Maximum Read Length (bp) | 128701 | 128655 |
| Minimum Read Length (bp) | 26 | 4 |
| Average Read Length (bp) | 6380 | 6345 |
| Median Read Length (bp) | 10549.5 | 21943 |
| Total Reads Length (bp) | 10425220236 | 10367243281 |
| Reads >= 100 bp | 1633670 | 1633203 |
| Reads >= 200 bp | 1625360 | 1619658 |
| Reads >= 500 bp | 1506349 | 1493906 |
| Reads >= 1 Kbp | 1339447 | 1328622 |
| Reads >= 10 Kbp | 344105 | 342604 |
| N50 value | 12732 | 12747 |
| Coverage | 22x |  |
