## Supplementary Table 9 for "Draft genome sequence of the pulse crop blackgram [*Vigna mungo* (L.) Hepper] reveals potential R-genes": Table S9 .docx

Table S9. Details of genes related to disease resistance, susceptibility and DNA damage repair identified in blackgram whole genome sequencing of cultivar PantU-31 through homology search with cowpea and mungbean

| **Protein ID** | **Protein Names** | **Gene Name** | **Organism Name** |
| --- | --- | --- | --- |
| g4.t1 | Disease resistance protein RPM1 | DEO72_LG10g4108 | Cowpea |
| g291.t1 | Nematode resistance protein-like HSPRO1 | DEO72_LG6g3289 | Cowpea |
| g980.t1 | nematode resistance protein-like HSPRO2 | LOC106765357 | Mung bean |
| g1162.t1 | putative disease resistance protein RGA4 isoform X1 | LOC106756847 | Mung bean |
| g1514.t1 | DNA damage-repair/toleration protein DRT100 | LOC106762878 | Mung bean |
| g1579.t1 | TIR domain-containing protein | Vigan.05G038400 VIGAN_05038400 | Vigna angularis var. angularis |
| g1908.t1 | probable disease resistance protein At5g66900 | LOC106768721 | Mung bean |
| g1909.t1 | probable disease resistance protein At5g66900 | LOC106755496 | Mung bean |
| g1910.t1 | probable disease resistance protein At5g66900 | LOC106755496 | Mung bean |
| g3904.t1 | probable disease resistance protein At5g66900 | LOC106777819 | Mung bean |
| g4787.t1 | LEAF RUST 10 DISEASE-RESISTANCE LOCUS RECEPTOR-LIKE PROTEIN KINASE-like 1.2 isoform X2 | LOC106771296 | Mung bean |
| g5186.t1 | nematode resistance protein-like HSPRO2 | LOC106757997 | Mung bean |
| g5379.t1 | DNA damage-repair/toleration protein DRT102 | LOC106774593 | Mung bean |
| g5778.t1 | DNA repair protein XRCC3 homolog | LOC106758811 | Mung bean |
| g5946.t1 | putative disease resistance RPP13-like protein 3 isoform X1 | LOC106779091 | Mung bean |
| g5947.t1 | disease resistance protein RPP13-like | LOC106756053 | Mung bean |
| g6384.t1 | LEAF RUST 10 DISEASE-RESISTANCE LOCUS RECEPTOR-LIKE PROTEIN KINASE-like 1.1 | LOC106771859 | Mung bean |
| g6385.t1 | LEAF RUST 10 DISEASE-RESISTANCE LOCUS RECEPTOR-LIKE PROTEIN KINASE-like 1.1 | LOC106770705 | Mung bean |
| g6683.t1 | protein ENHANCED DOWNY MILDEW 2 | LOC106775877 | Mung bean |
| g7021.t1 | putative disease resistance protein At1g50180 | LOC106759875 | Mung bean |
| g7141.t1 | disease resistance-like protein CSA1 | LOC106763560 | Mung bean |
| g7231.t1 | disease resistance protein RPP8 | LOC106777977 | Mung bean |
| g7410.t1 | TIR domain-containing protein (Fragment) | PHAVU_010G0278000g | Phaseolus vulgaris |
| g7538.t1 | probable disease resistance protein At4g19060 | LOC106775463 | Mung bean |
| g7944.t1 | disease resistance protein TAO1 isoform X2 | LOC106753259 | Mung bean |
| g8592.t1 | DNA mismatch repair protein | LR48_Vigan03g279200 | Phaseolus angularis |
| g8869.t1 | DNA damage-repair/toleration protein DRT100 | LOC106759210 | Mung bean |
| g8930.t1 | disease resistance protein RPM1 | LOC106777972 | Mung bean |
| g9212.t1 | pathogen-related protein | LOC106778131 | Mung bean |
| g9496.t1 | DNA mismatch repair protein PMS1 isoform X1 | LOC106769770 | Mung bean |
| g10184.t1 | protein ENHANCED PSEUDOMONAS SUSCEPTIBILTY 1-like | LOC106766259 | Mung bean |
| g10466.t1 | double-strand break repair protein MRE11 isoform X1 | LOC106780307 | Mung bean |
| g10566.t1 | TMV resistance protein N isoform X1 | LOC106761051 | Mung bean |
| g10604.t1 | DNA damage-repair/toleration protein DRT100 | LOC106759768 | Mung bean |
| g10985.t1 | pathogenesis-related protein 2-like | LOC106773581 | Mung bean |
| g10986.t1 | pathogenesis-related protein 2-like | LOC106773664 | Mung bean |
| g10988.t1 | pathogenesis-related protein 2-like | LOC106773664 | Mung bean |
| g10989.t1 | pathogenesis-related protein 2-like | LOC106773581 | Mung bean |
| g10990.t1 | pathogenesis-related protein 2-like | LOC106773581 | Mung bean |
| g10991.t1 | pathogenesis-related protein 2-like | LOC106773581 | Mung bean |
| g11326.t1 | DNA excision repair protein ERCC-4 | DEO72_LG11g2839 | Cowpea |
| g11722.t1 | DNA repair protein RAD5A | LOC106758090 | Mung bean |
| g11871.t1 | LEAF RUST 10 DISEASE-RESISTANCE LOCUS RECEPTOR-LIKE PROTEIN KINASE-like 1.4 isoform X3 | LOC106758525 | Mung bean |
| g11997.t1 | probable disease resistance protein At4g33300 | LOC106765225 | Mung bean |
| g11998.t1 | probable disease resistance protein At4g33300 | LOC106764928 | Mung bean |
| g12142.t1 | TMV resistance protein N-like | LOC106763241 | Mung bean |
| g12187.t1 | TMV resistance protein N-like isoform X1 | LOC106757003 | Mung bean |
| g12631.t1 | TMV resistance protein N (Fragment) | N CR513_35863 | Mucuna pruriens |
| g13079.t1 | pathogenesis-related protein 1C-like | LOC106770364 | Mung bean |
| g13134.t1 | DNA mismatch repair protein MutS2 | DEO72_LG2g4696 | Cowpea |
| g13267.t1 | DNA mismatch repair protein MSH3 | LOC106769933 | Mung bean |
| g13331.t1 | DNA repair protein REV1 | DEO72_LG11g3249 | Cowpea |
| g13332.t1 | DNA repair protein REV1 | DEO72_LG11g3249 | Cowpea |
| g13587.t1 | DNA damage-inducible protein 1 | LOC106774443 | Mung bean |
| g14283.t1 | disease resistance protein At4g27190 | LOC106754308 | Mung bean |
| g14543.t1 | rust resistance kinase Lr10-like | LOC106755174 | Mung bean |
| g14544.t1 | LEAF RUST 10 DISEASE-RESISTANCE LOCUS RECEPTOR-LIKE PROTEIN KINASE-like 2.2 | LOC106755211 | Mung bean |
| g14632.t1 | DNA repair protein RAD5B isoform X1 | LOC106769931 | Mung bean |
| g14659.t1 | putative disease resistance RPP13-like protein 1 | LOC106769968 | Mung bean |
| g14894.t1 | DNA mismatch repair protein MLH1 isoform X5 | LOC106762345 | Mung bean |
| g15628.t1 | LEAF RUST 10 DISEASE-RESISTANCE LOCUS RECEPTOR-LIKE PROTEIN KINASE-like 1.5 | LOC106777701 | Mung bean |
| g15918.t1 | putative disease resistance protein RGA1 | LOC106776707 | Mung bean |
| g16165.t1 | protein ENHANCED DISEASE RESISTANCE 2 | LOC106770618 | Mung bean |
| g16444.t1 | DNA damage-repair/toleration protein DRT100 | LOC106755598 | Mung bean |
| g16597.t1 | disease resistance protein RGA2-like | LOC106769976 | Mung bean |
| g16958.t1 | protein PLANT CADMIUM RESISTANCE 7 | LOC106767114 | Mung bean |
| g17020.t1 | LEAF RUST 10 DISEASE-RESISTANCE LOCUS RECEPTOR-LIKE PROTEIN KINASE-like 1.2 | LOC106766190 | Mung bean |
| g17266.t1 | protein ENHANCED DISEASE RESISTANCE 2-like | LOC106760089 | Mung bean |
| g17378.t1 | DNA repair protein XRCC3 homolog | LOC106758811 | Mung bean |
| g18093.t1 | disease resistance protein TAO1 isoform X2 | LOC106753259 | Mung bean |
| g18164.t1 | pathogenesis-related protein 5-like | LOC106763779 | Mung bean |
